# Multi-omics analysis and genome-scale metabolic reconstruction of cattle *Bos taurus* for optimal production of cultured meat

**DOI:** 10.1101/2024.12.09.627468

**Authors:** Junkyu Lee, Jaehyun Kim, Hyun Won Bae, Minji Kim, Byung Kwon Jung, Juyeon Kim, SungGun Lee, Hyun Uk Kim

## Abstract

With the growing urgency of addressing climate change, cultured meat has gained significant attention as a sustainable alternative to conventional meat production. *Bos taurus*, a key cattle species, is considered as a potential source of cultured meat. However, much remains to be understood about the biology of *B. taurus* muscle cells. In this study, bovine satellite cells (BSCs) derived from the semimembranosus muscle of Korean Hanwoo cattle were subjected to multi-omics profiling and genome-scale metabolic reconstruction. First, differential gene expression and gene set enrichment analyses, based on RNA-seq data, identified key pathways associated with muscle cell proliferation (e.g., ‘Cell cycle’ and ‘RNA polymerase’) and differentiation (e.g., ‘Cytoskeleton in muscle cells’ and ‘Tryptophan metabolism’). Next, using the human1 GEM as a template, we constructed the first *B. taurus*-specific genome-scale metabolic model (GEM), named BtaSBML2986, which comprises 2,986 genes, 13,278 reactions, and 8,652 metabolites. Muscle cells were cultured under six distinct conditions, and biomass predictions generated using BtaSBML2986 were validated against experimental growth rates. This integrated approach also provided insights into core pathways such as glycolysis and the TCA cycle. BtaSBML2986 represents a significant step forward in understanding *B. taurus* muscle metabolism and will serve as a valuable tool for advancing cultured meat research and optimizing culture processes.

## Introduction

Cultured meat, produced by cultivating animal cells in vitro, offers a sustainable solution to the environmental and ethical challenges of traditional livestock farming (Bryant, 2020). Compared to conventional meat production, it requires significantly less land and water, produces fewer greenhouse gas emissions, and eliminates the need for animal slaughter, addressing both environmental and animal welfare concerns (Treich, 2021). As global demand for meat rises, cultured meat presents a viable alternative for meeting consumer needs while mitigating environmental impacts. Optimizing muscle cell growth and metabolism is critical to the success of cultured meat production, particularly for species such as *Bos taurus*, a key source of global meat supply.

Recent advances in omics technologies (e.g., including genomics, transcriptomics, proteomics, and metabolomics) have significantly enhanced our understanding of biological systems (Ramalhete et al., 2023; Sonehara & Okada, 2021; Zou et al., 2021). These tools have been increasingly applied to livestock science, enabling researchers to unravel the molecular mechanisms underlying growth, development, and production traits in economically important species such as *B. taurus* (Yu et al., 2023). Studies have employed transcriptomics to identify genes involved in muscle growth and differentiation (Deng et al., 2024), while metabolomics has provided insights into metabolic profiles influencing meat flavor, tenderness, and nutritional value (Carrillo et al., 2016). However, these approaches often lack integration, limiting their ability to fully capture the complexity of bovine muscle metabolism.

Genome-scale metabolic models (GEMs) are powerful tools for integrating multi-omics data, providing a systems-level representation of an organism’s metabolism (Gu et al., 2019). Such models enable the identification of key pathways driving muscle growth, the simulation of metabolic responses under varying conditions, and the design of optimal culture media to support cell proliferation and differentiation. In the context of cultured meat, a B. taurus-specific GEM could significantly improve production efficiency while reducing costs, making cultured meat more accessible and sustainable. Despite their potential, GEMs specific to *B. taurus* muscle cells are lacking. Existing models often rely on extrapolated data from other species, failing to capture the unique metabolic characteristics of bovine muscle tissue (Wang et al., 2021). This gap hinders system-level analyses and limits advancements in both traditional meat production and cultured meat research. Therefore, the development of a B. taurus-specific GEM is critical for addressing these challenges.

This study aims to investigate the gene expression profiles of *B. taurus* muscle cells using RNA sequencing and reconstruct their GEM, BtaSBML2986, which was validated with transcriptomic and metabolomic data derived from Hanwoo cattle. Differential gene expression and gene set enrichment analyses (GSEA) were performed to identify key pathways driving muscle proliferation and differentiation, revealing distinct metabolic and regulatory features at various growth stages. The reconstructed GEM, BtaSBML2986, was validated through growth rate predictions and flux analysis of glycolysis and the TCA cycle, highlighting its utility in simulating metabolic phenotypes under varying conditions. By combining RNA-seq data with GEM simulations, this study offers insights for understanding *B. taurus* muscle metabolism, and improving traditional livestock production.

## Results and discussion

### Cultivation of the *B. taurus* cells

In this study, bovine satellite cells (BSCs) derived from the semimembranosus muscle of Korean Hanwoo cattle were used for omics analysis and genome-scale metabolic reconstruction. BSCs were prepared from three individual cattle, designated as ‘No. 43,’ ‘No. 45,’ and ‘No. 66’, and cultured in three different media: ‘885’ (relatively low glucose, 1.0 g/L, and sodium pyruvate, 110.0 mg/L), ‘965’ (relatively high glucose, 4.5 g/L), and ‘995’ (relatively high glucose, 4.5 g/L, and sodium pyruvate, 110.0 mg/L) (Table 1). These media, based on Dulbecco’s Modified Eagle Medium (DMEM), varied in carbon content and were prepared in two formulations, with 10% fetal bovine serum (FBS) to promote proliferation and 3% horse serum (HS) to induce differentiation, resulting in six distinct media conditions. Cells were sampled at 24 hours and 72 hours during cultivation, with samples grown in media containing 10% FBS designated as ‘P1’ (24 h) and ‘P3’ (72 h) to indicate proliferation, and those grown in media with 3% HS designated as ‘D1’ (24 h) and ‘D3’ (72 h) to indicate differentiation. This experimental setup allowed the generation of a comprehensive dataset across three cell sample sets, six media conditions, and two distinct growth and differentiation stages.

**Table 1.** Definition of the cell sample sets and the cultivation conditions.

| Cell sample sets from *Bos taurus* | 43 | 45 | 66 |
| --- | --- | --- | --- |
| **Medium (Proliferation)** | 885 (DMEM with 1.0 g/L glucose and 110.0 mg/L sodium pyruvate; Gibco, 11885084), 10% FBS | 965 (DMEM with 4.5 g/L glucose; Gibco, 11965175), 10% FBS | 995 (DMEM with 4.5 g/L glucose and 110.0 mg/L sodium pyruvate; Gibco, 11995115), 10% FBS |
| **Medium (Differentiation)** | 885 (DMEM with 1.0 g/L glucose and 110.0 mg/L sodium pyruvate; Gibco, 11885084), 3% HS | 965 (DMEM with 4.5 g/L glucose; Gibco, 11965175), 3% HS (Gibco, 26050070) | 995 (DMEM with 4.5 g/L glucose and 110.0 mg/L sodium pyruvate; Gibco, 11995115), 3% HS (Gibco, 26050070) |

\*DMEM, Dulbecco’s Modified Eagle Medium; FBS, fetal bovine serum (Cytiva, SH883IH2540); HS, horse serum (Gibco, 26050070)

### Transcriptome analysis of the *B. taurus* cells

Transcriptomic analysis of 72 samples revealed distinct clustering patterns based on sampling points (P1, P3, D1, and D3), as visualized in the UMAP plots (Fig. 1A). This result indicates that transcriptome profiles are predominantly influenced by sampling points, irrespective of the medium or cell sample sets. Using this insight, we performed GSEA to identify pathways significantly enriched between the P1 and P3, as well as D1 and D3 sampling points. Conditions for each sample were assigned based on its respective sampling point, as illustrated in the comparative analyses (Figs. 1B and 1C).

**Fig. 1.**
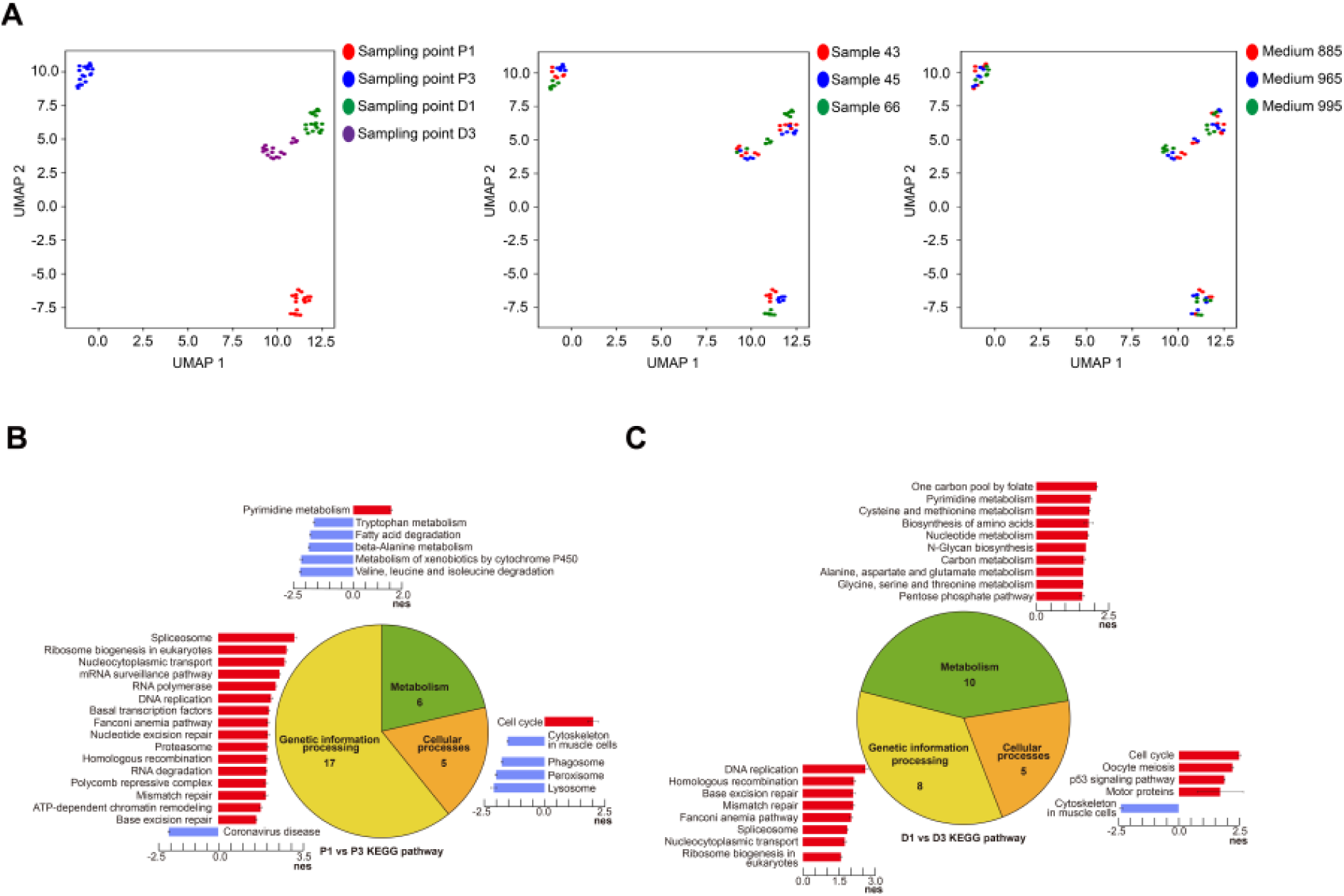
Transcriptomic analysis of bovine satellite cells (BSCs) derived from *B. taurus*. Transcriptome data were obtained twice from four different sampling points (P1 versus P3, and D1 versus D3), using three cell sample sets, and three media, resulting in a total of 72 data points. (A) UMAP plots for the three different conditions: sampling points, cell sample sets, and media. (B) and (C) Gene set enrichment analysis between cell samples collected at P1 and P3, and at D1 and D3, revealing significantly enriched pathways (P-value < 0.05, FDR < 0.05, and |NES| > 1.5). In the bar graph, positive values indicate enrichment in P1 and D1 in comparison with P3 and D3, respectively, while negative values indicate the opposite, which enrichment in P3 and D3.

In the comparison between P1 and P3, pathways such as ‘Spliceosome,’ ‘Ribosome biogenesis in eukaryotes,’ ‘RNA polymerase,’ and ‘DNA replication’ were overexpressed at P1, highlighting heightened genetic information processing. Additionally, pathways related to metabolism and cellular processes, such as ‘Pyrimidine metabolism’ and ‘Cell cycle,’ were also enriched at P1. These findings suggest active transcription, translation, and cell division at the P1 stage, indicative of rapid growth and proliferation. Conversely, at P3, pathways such as ‘Coronavirus disease,’ ‘Tryptophan metabolism,’ ‘Fatty acid degradation,’ ‘Valine, leucine, and isoleucine degradation,’ ‘Cytoskeleton in muscle cells,’ ‘Peroxisome,’ and ‘Lysosome’ were enriched. These pathways are associated with metabolic flexibility, immune response, and structural muscle cell components, reflecting greater differentiation and maturity at the P3 stage.

Similarly, in the comparison between D1 and D3, most pathways were significantly enriched in D1, with the exception of the ‘Cytoskeleton in muscle cells’ pathway, which was enriched at D3. At D1, metabolic pathways such as ‘Biosynthesis of amino acids,’ ‘Nucleotide metabolism,’ and ‘Carbon metabolism’ were prominent, underscoring rapid cell growth and division. Additionally, pathways related to genetic information processing, including ‘DNA replication’ and ‘DNA repair,’ suggest active DNA synthesis and maintenance. The enrichment of the ‘Cell cycle’ and ‘p53 signaling pathway’ at D1 further supports the stage’s emphasis on cell proliferation and regulation. In contrast, the D3-specific enrichment of the ‘Cytoskeleton in muscle cells’ pathway emphasizes structural muscle development, including sarcomere formation and the organization of actin and myosin filaments critical for muscle contraction. Together, these findings indicate that D1 samples exhibit robust proliferative activity, whereas D3 samples display characteristics of structural specialization and differentiation.

To further explore the biological differences between sampling points, we analyzed log2FoldChange values of representative genes associated with key pathways involved in proliferation and differentiation (Fig. 2). In the comparison between P1 and P3 (Fig. 2A), pathways such as ‘Cell cycle,’ ‘Ribosome biogenesis in eukaryotes,’ ‘Pyrimidine metabolism,’ ‘DNA replication,’ and ‘RNA polymerase’ were examined. For each pathway, the top five genes with the highest expression levels in P1 and P3 were identified. The findings reveal that P1 samples exhibit characteristics of an early, active proliferation stage, with elevated expression of genes involved in cell cycle progression, RNA synthesis, and DNA replication. For example, *MYC*, which promotes cell cycle progression, and *POLR3G*, critical for RNA synthesis during early proliferation, were highly expressed in P1. This reflects an increased demand for transcriptional and translational machinery, underscoring rapid growth and proliferation. In contrast, P3 samples displayed a distinct focus on cellular differentiation and maturation. Genes such as *CDKN1C*, which inhibits cell cycle progression to facilitate differentiation, and *POLR3GL*, which maintains RNA synthesis in more mature cells, were more highly expressed. Additionally, genes such as *UTP14A*, associated with ribosome assembly, and *DPYD*, involved in pyrimidine metabolism, were enriched in P3, indicating an emphasis on cellular maintenance and structural specialization. Together, these results highlight a transition from the proliferative, early-growth characteristics of P1 to the differentiated and mature state of P3.

**Fig. 2.**
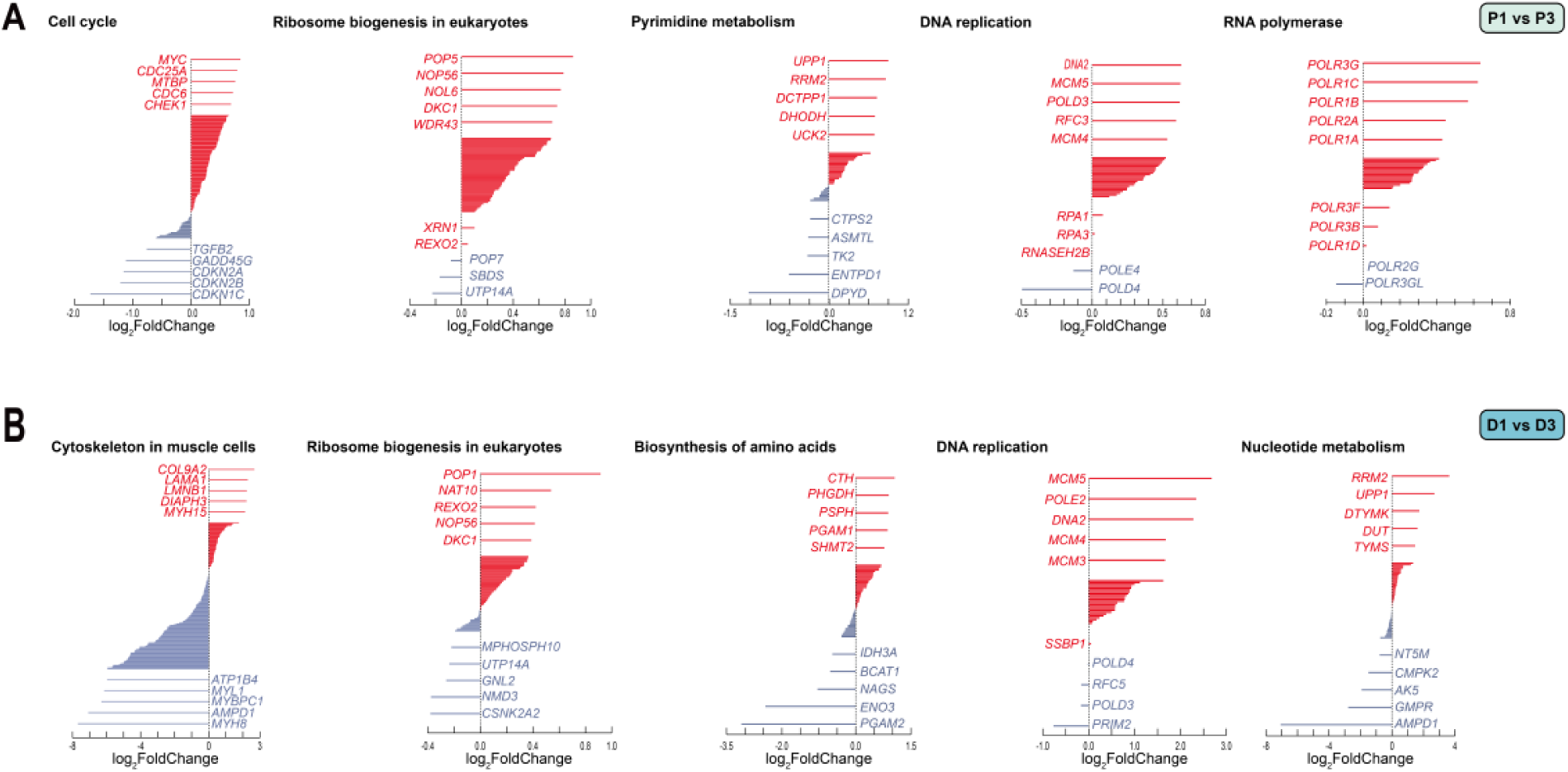
Representative genes associated with significantly enriched pathways, obtained through gene set enrichment analysis, and changes in their corresponding expression values (log2FoldChange). (A) The top five genes with the highest fold change values between P1 and P3 for each of the significantly enriched pathways presented. (B) The top five genes with the highest fold change values between D1 and D3 for each of the significantly enriched pathways presented.

Similarly, the comparison between D1 and D3 (Fig. 2B) identified key genes associated with pathways such as ‘Cytoskeleton in muscle cells,’ ‘Ribosome biogenesis in eukaryotes,’ ‘Biosynthesis of amino acids,’ ‘DNA replication,’ and ‘Nucleotide metabolism.’ The analysis revealed that D1 samples are primarily focused on active proliferation, with high expression of genes such as *MCM5*, *PRIM2*, and *RRM2*, which are essential for DNA replication and nucleotide synthesis. Genes such as *POP1* and *CSNK2A2*, which contribute to ribosome biogenesis, further support the production of ribosomes necessary for protein synthesis during this stage. By contrast, D3 samples were enriched with genes associated with muscle differentiation and energy metabolism. For instance, *MYH8*, a key gene in muscle fiber development and regeneration, showed significantly higher expression in D3, reflecting advanced muscle differentiation. Similarly, *PGAM2*, crucial for energy metabolism in muscle tissue, and *AMPD1*, which aids in ATP regeneration during recovery, were more highly expressed in D3. The enrichment of *CTH*, involved in cysteine metabolism and antioxidant defense, further underscores D3’s focus on metabolic adaptation and energy optimization. These findings indicate a shift from the proliferative activities of D1 to the differentiated and functional muscle state of D3.

### Genome-scale metabolic reconstruction of *B. taurus*

The *B. taurus* GEM was constructed using a systematic workflow that integrates biological data from ARS-UCD2.0 at NCBI (Sayers et al., 2022), KEGG (Kanehisa & Goto, 2000), and Rhea (Bansal et al., 2022) (Fig. 3). ARS-UCD2.0 is a genome assembly version for *B. taurus* available at NCBI. Human1, a highly curated human GEM, served as the template model. Genes and their associated protein sequences were extracted from Human1 and annotated with UniProt IDs to perform ortholog analysis. Using the ARS-UCD2.0, bidirectional BLASTP was performed with DIAMOND (Buchfink et al., 2021) to map *B. taurus* genes to their human orthologs, resulting in an ortholog GEM. To incorporate species-specific reactions and metabolites, metabolic data linked to *B. taurus* were extracted from KEGG and Rhea, with additional refinement based on information from Pathway Tools (Karp et al., 2021) and the literature. Rhea data were prioritized to enhance the model’s robustness, addressing limitations of KEGG. A draft GEM was constructed by integrating the ortholog GEM and species-specific metabolic network data, and GPRuler was used to refine gene-protein-reaction (GPR) associations. This iterative process resulted in the development of the finalized *B. taurus* GEM, BtaSBML2986.

**Fig. 3.**
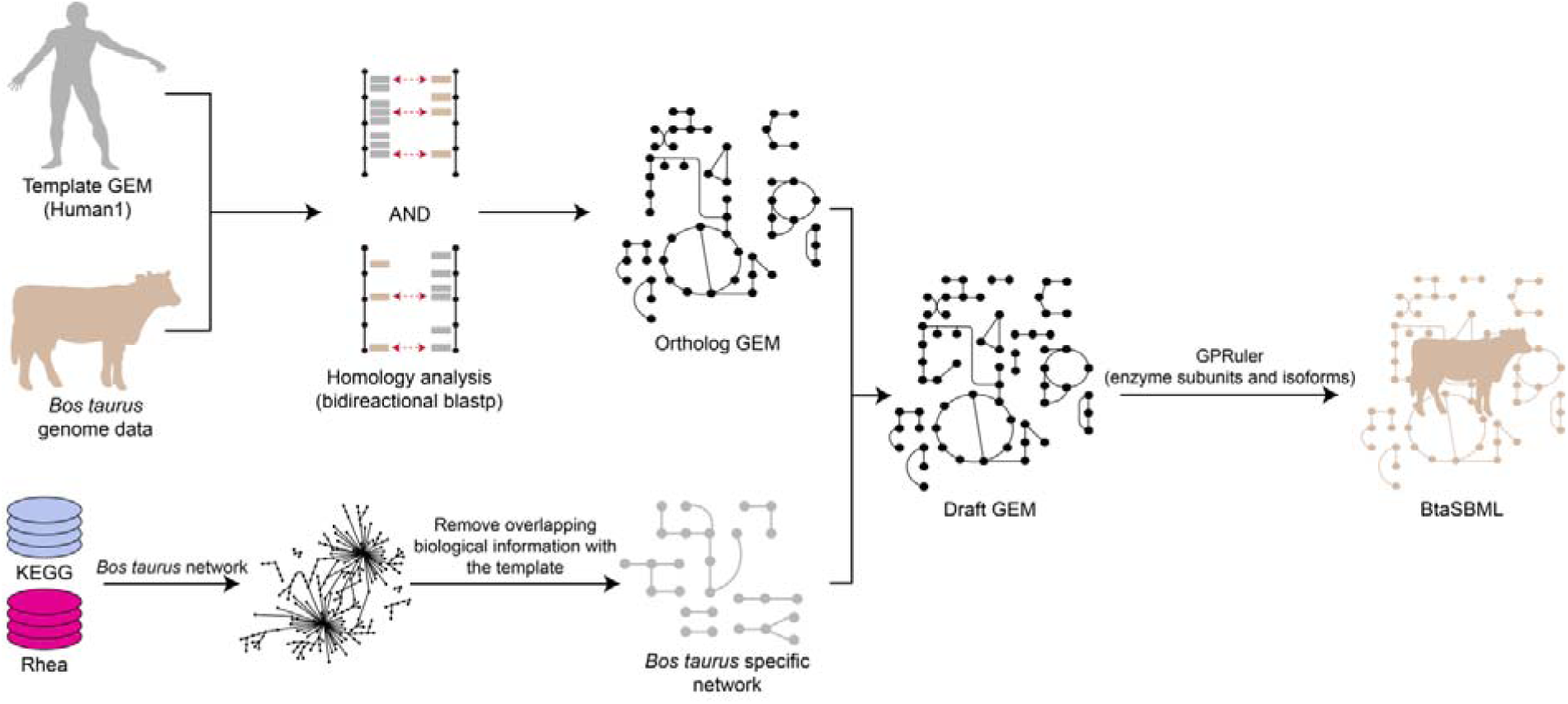
A workflow for the generation of a genome-scale metabolic model of *B. taurus*. First, an ortholog GEM was generated by directional homology analysis of the enzymes that comprise Human1 (Robinson et al., 2020), a reference human GEM, and *B. taurus* genome. Second, *B. taurus* metabolic network was generated by extracting *B. taurus*-specific (denoted by the organism code for *Bos taurus*) information from KEGG and Rhea databases. Next, a draft GEM of *B. taurus* was generated by integrating the ortholog GEM and the *B. taurus*-specific metabolic network. Finally, BtaSBML2986, the final *Bos taurus* GEM, was generated by refining the draft GEM for mass and charge balances, and modifying its gene-protein-reaction (GPR) associations using GPRuler (Di Filippo et al., 2021).

The resulting *B. taurus* GEM, BtaSBML2986, comprises 2,986 genes, 13,278 reactions, and 8,652 metabolites (Fig. 4C). Model quality was assessed using MEMOTE, achieving a total score of 79% and a consistency score of 88%, a key metric representing the model’s feasibility. This consistency score matches that of Human1, underscoring the high quality of BtaSBML2986 (Fig. 4D). Notably, eight new subsystems were introduced into BtaSBML2986 that are absent in Human1 and iCHOv1 (Hefzi et al., 2016), the GEM of Chinese hamster ovary (CHO) cells. These subsystems include ‘Biosynthesis of cofactors,’ ‘Carbapenem biosynthesis,’ ‘Chlorocyclohexane and chlorobenzene degradation,’ ‘Fluorobenzoate degradation,’ ‘Mannose type O-glycan biosynthesis,’ ‘Selenocompound metabolism,’ ‘Toluene degradation,’ and ‘alpha-linolenic acid metabolism’ (Fig. 4A). Among these, steroid metabolism exhibited the largest number of added reactions, followed by ‘Mannose type O-glycan biosynthesis,’ ‘Folate metabolism,’ and ‘Chlorocyclohexane and chlorobenzene degradation’. Finally, comparative analysis of essential genes using COBRApy (Ebrahim et al., 2013) revealed 17 essential genes in BtaSBML2986, 91 in Human1, and 37 in iCHOv1 (Fig. 4B). These findings highlight the unique metabolic features of BtaSBML2986 that require further studies.

**Fig. 4.**
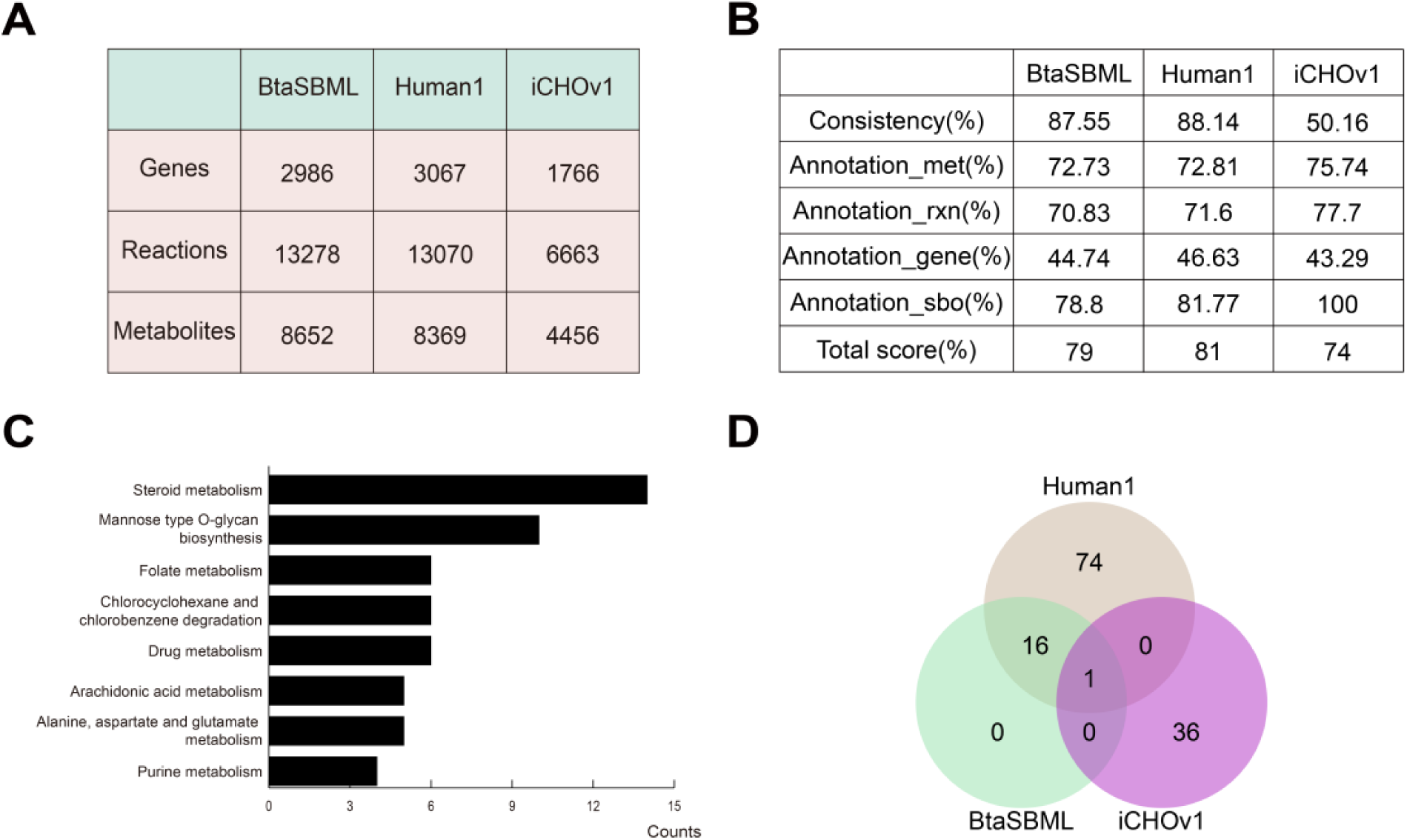
Features of the B. taurus GEM, BtaSBML2986. (A) Table summarizing the number of genes, reactions, and metabolites in each model. (B) Results of the MEMOTE applied to each model. (C) Bar plot representing the number of reactions in the 8 newly introduced subsystems in BtaSBML2986, which are not available in Human 1 (Robinson et al., 2020), the reference human GEM, and iCHOv1 (Hefzi et al., 2016), the GEM of Chinese hamster ovary (CHO) cells. (D) Venn diagram illustrating the number of essential genes shared among the three GEMs. Essential genes were predicted using COBRApy.

### Experimental validation of BtaSBML2986

To validate the *B. taurus* GEM, BtaSBML2986, we first compared its predictions with experimental data for growth rates and differentially abundant metabolites (Fig. 5). Experimental growth rate data were obtained for three cell sample sets (‘43,’ ‘45,’ and ‘66’) under three media conditions (‘885,’ ‘965,’ and ‘995’). For these validation simulations, cell-specific GEMs were first reconstructed by integrating BtaSBML2986 with RNA-seq data for each condition. For the comparison with the experimental growth data, flux balance analysis (FBA), constrained by uptake rates of major nutrients derived from media composition analysis, was performed using biomass reaction as the objective function to predict growth rates. The predicted growth rates closely matched the experimentally observed values (Fig. 5A). Next, metabolome data were used to validate the model by comparing metabolites with significantly different flux-sum (Kim et al., 2007) values, predicted using P1- and P3-specific GEMs, to those with significant concentration differences identified through metabolome analysis. Here, flux-sum is a total sum of all the fluxes associated with the consumption or generation of a metabolite, and indicates the biological importance of a metabolite (Lee et al., 2024). Flux-sum values of entire metabolites were calculated by simulating the cell-specific GEMs using least absolute deviation (LAD) (Lee et al., 2022; Kim et al., 2011). LAD was used to calculate reaction fluxes as close possible as reference flux weights, calculated from RNA-seq, thereby generating reliable genome-scale flux distributions of each cell. When performing LAD, the biomass reaction was set as the objective function, and constraints were provided, based on medium composition and experimental growth rate data. As a result, significant differences in both datasets were assessed using the Mann-Whitney U test with a Benjamini-Hochberg (BH)-corrected P-value < 0.05 (Fig. 5B). These results demonstrate the utility of BtaSBML2986 in accurately predicting growth rates and identifying differentially abundant metabolites under varying conditions.

**Fig. 5.**
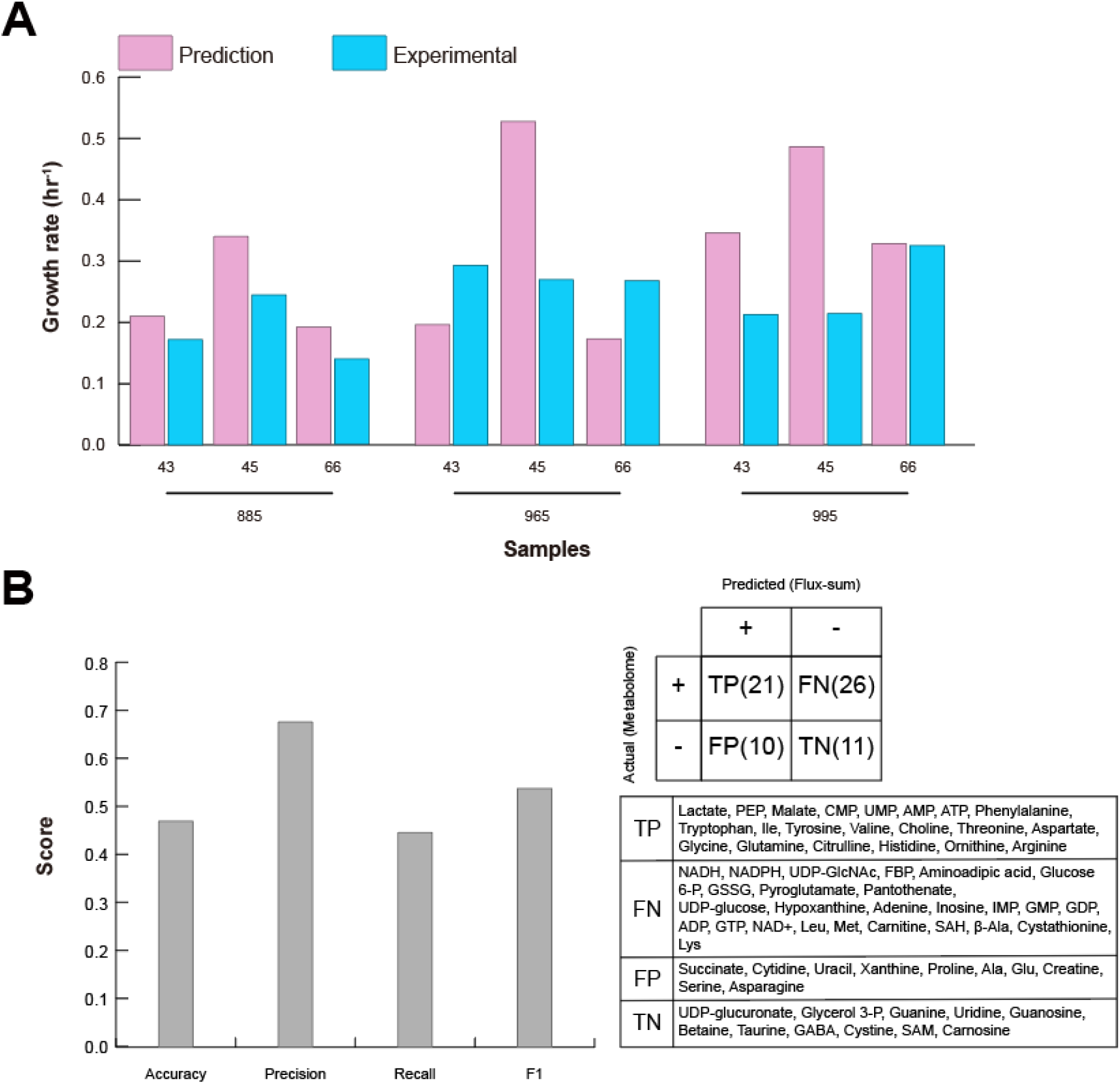
Validation of the *B. taurus* GEM, BtaSBML2986. (A) Comparison of the experimentally obtained growth rates and those predicted using BtaSBML2986 for the three cell sample sets (i.e., ‘43’, ‘45’, and ‘66’) and three media conditions (i.e., ‘885’, ‘965’, and ‘995’). (B) Evaluation of the predictions of differentially abundant metabolites between the P1 and P3 samples. Metabolites with significantly different flux-sum values were predicted using P1 and P3 specific GEMs, which were generated using BtaSBML2986. They were next compared with metabolites with significantly different concentrations between P1 and P3, which were obtained through metabolome analysis performed in this study. Metabolites with significantly different flux-sum values using the GEMs or those with significantly different concentrations P1 and P3 were identified using Mann-Whitney U test (BH-corrected P-value < 0.05).

### Flux analysis of glycolysis and TCA cycle using the cell-specific GEMs

Glycolysis and TCA cycle in the *B. taurus* cells at P1 and P3 were further examined by using the cell-specific GEMs across the three media as analyzing these pathways offer valuable insights into the production of essential metabolites and their impact on biomass generation and cellular growth (Fig. 6). For this, the flux distributions of the nine P1- and P3-specific GEMs predicted using LAD mentioned above (Fig. 5B) were used to analyze their fluxes through glycolysis and TCA cycle; samples ‘No_43_P3_965’ and ‘No_66_P3_965’ were excluded due to infeasible solutions. As a result, the analysis revealed active glycolysis, indicated by positive fluxes from glucose to acetyl-CoA, demonstrating its role in ATP production, fatty acid biosynthesis, and amino acid metabolism. Fluxes toward citrate and succinate confirmed TCA cycle activity, supporting oxidative phosphorylation and the biosynthesis of fatty acids and sterols through citrate export to the cytosol. Notably, while the flux from oxalosuccinate to alpha-ketoglutarate (AKG) was reduced to zero during optimization, LAD-based predictions showed that AKG biosynthesis was maintained via the isocitrate-to-AKG pathway. This highlights the model’s flexibility in sustaining essential metabolite production through alternative routes when specific reactions are deprioritized during optimization.

**Fig. 6.**
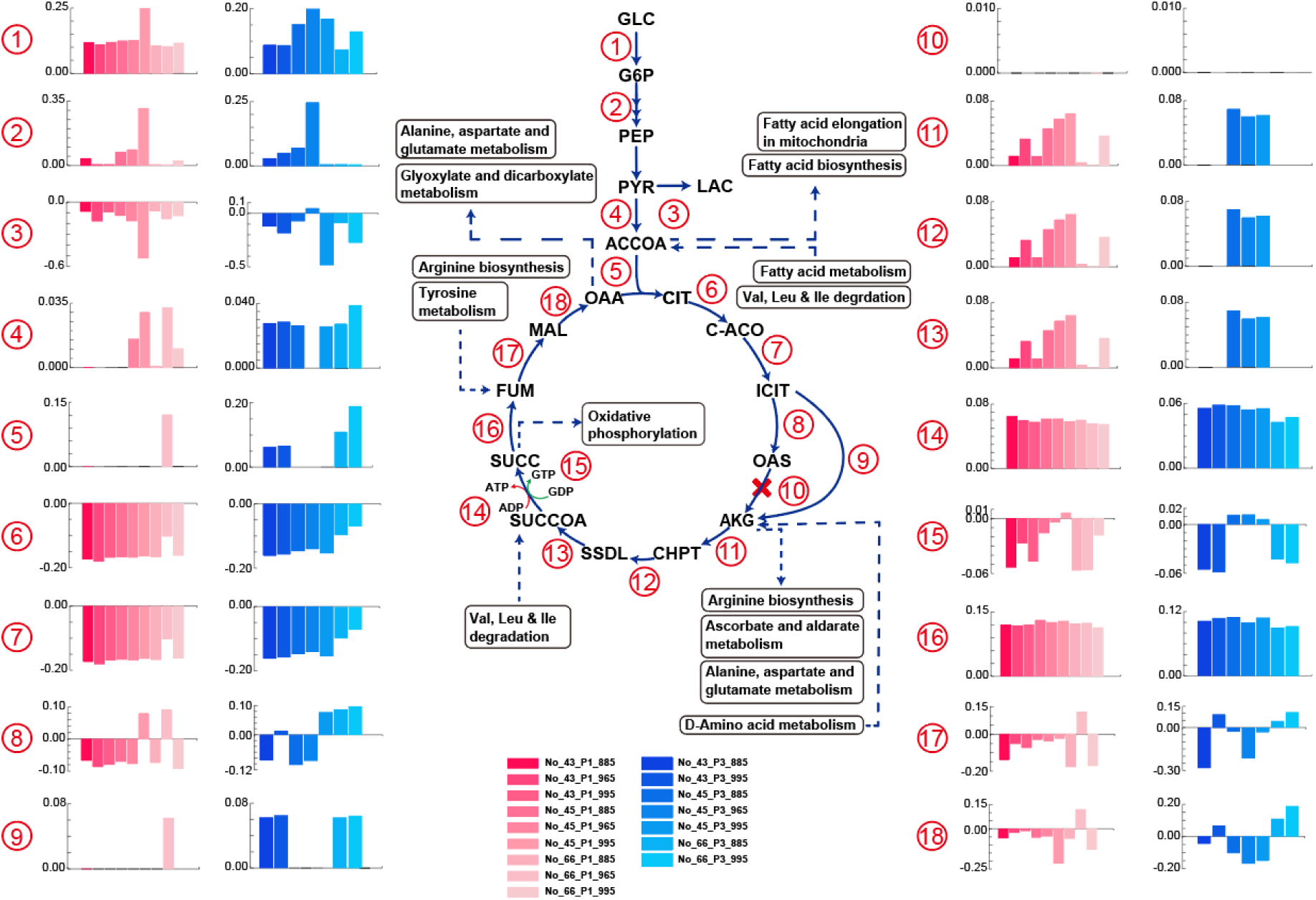
Flux analysis of glycolysis and TCA cycle for the three cell sample sets at P1 and P3 time points across the three media. The presented red and blue bar plots indicate flux data obtained for the P1 and P3 conditions, respectively. Numbers on the left side of each bar graph correspond to the flux of the reaction with the same number in glycolysis and TCA cycle. To simplify visualization, all flux directions are presented in one direction, with negative values in the bar graph indicating flux in the opposite direction. Abbreviations: GLC: D-glucose, G6P: glucose-6-phosphate, PEP: phosphoenol-pyruvate, PYR: pyruvate, LAC: lactate, ACCOA: acetyl-CoA, CIT: citrate, C-ACO: cis-aconitate, ICIT: isocitrate, OAS: oxalosuccinate, AKG: alpha-ketoglutaric acid, CHPT: 3-carboxy-1-hydroxypropyl-ThPP, SSDL: S-succinyl-dihydrolipoamide-E, SUCCOA: succinyl-CoA, SUCC: succinate, FUM: fumarate, MAL: (S)-malate, OAA: oxaloacetate.

## Materials and methods

### Cell cultivation

BSCs from the Korean Hanwoo cattle, ‘No. 43,’ ‘No. 45,’ and ‘No. 66’, were cultured under the six different media (Table 1) as follows. For each culture medium, BSCs were seeded at 1×10^4^ cells per well in a 6-well plate to initiate proliferation. Cell sampling for transcriptome and metabolome analysis, as well as cell counting, were conducted at 1 day (24 h) and 3 days (72 h) after the start of proliferation. On the fourth day of proliferation (96 h), the medium was replaced with a differentiation medium to initiate differentiation. During differentiation, cell sampling for transcriptome analysis was performed at 1 day (24 h) and 3 days (72 h). Cell counts were measured using the Cellometer K2 device (Nexcelom Bioscience).

### RNA sequencing

In case of transcriptome data, total RNA was extracted from semimembranous muscle using the TRIzol reagent (Invitrogen) following the manufacturer’s instructions. RNA samples were treated with DNase1 (0.1%; DNA-Free, Ambion) to remove DNA contaminants, and RNA quality and quantity were assessed using a Bioanalyzer 2100 with RNA 6000 Nano Labchips (Agilent Technologies). Only samples with an RNA integrity number (RIN) ≥ 7 were used for further analysis. Complementary DNA (cDNA) libraries were synthesized using the Illumina TruSeq kit (Illumina) following the manufacturer’s instructions, and sequencing was performed on the NovaSeq 6000 platform (Illumina), generating more than 60 million paired-end reads (2 × 101 bp). For RNA-seq data analysis, sequencing adapters and low-quality bases were trimmed using Trimmomatic 0.39, retaining reads with a Phred quality score >15 in a 4-base wide sliding window and read lengths >36 bp. The Hanwoo genome was constructed using DNA sequencing data from 21 Hanwoo samples provided by the National Institute of Animal Science (NIAS), Korea, and the *Bos taurus* UMD 3.1.1 genome. Low-quality bases (Phred score < 20) were removed using Cutadapt 2.1, and reads longer than 50 bp were aligned to the Bos taurus UMD 3.1.1 genome using BWA 0.7.17. Hanwoo-specific sequences were identified using GATK 4.2.4.1, including SortSam, MarkDuplicates, and HaplotypeCaller, and the Hanwoo genome sequence was generated using in-house python scripts and gVCF files. Transcript expression values were quantified using STAR 2.7.10b and RSEM 1.2.28 with the Hanwoo genome as the reference. We utilized expected_count and transcripts per million (TPM) values, considering transcripts with expected_count ≥ 5 and TPM ≥ 0.3 across all replicates as expressed transcripts.

### RNA-seq data analysis

Before conducting UMAP, the RNA-seq data contained separate expression values for isoforms, so the expression values of all isoforms for each gene were summed using TPM values to ensure accurate gene-level expression. UMAP was then implemented using the umap-learn package (McInnes et al., 2018) with default settings, including n_neighbors set to 15, which determines the number of neighboring points used in the local approximation of manifold structure. Larger values for n_neighbors preserve more global structure but may reduce detailed local structure. The min_dist parameter was set to the default value of 0.1, controlling how tightly the embedding is allowed to compress points together, striking a balance between local accuracy and even distribution of embedded points. The random_state was fixed at 42 to ensure reproducibility. Differential gene expression analysis and GSEA were performed using the pyDESeq2 (Muzellec et al., 2023) package. To create a *B. taurus*-specific gene set, gene sets specific to *B. taurus* were extracted from KEGG and used as KEGG_2024_Cow.gmt. Using this customized gene set, GSEA was conducted on nine samples each from the P1, P3, D1, and D3 groups using the pyDESeq2 package. Significantly enriched pathways were identified based on thresholds of P*-*value < 0.05, false discovery rate (FDR) < 0.05, and absolute normalized enrichment score (|NES|) > 1.5.

### Metabolome analysis

Before analyzing the metabolome sample data, a preprocessing step was conducted. Intracellular metabolites were sampled using methanol quenching followed by rapid filtration. Metabolite extraction was performed using a modified Folch extraction method, which employs a biphasic mixture of chloroform, methanol, and water (2.5:2.5:1) as described by (Jazmin et al., 2017). To analyze metabolome samples, the following steps were conducted. LC-MS analysis for intracellular amino acids were performed on a Waters Acquity UPLC I-class & Xevo G2-XS QTof system equipped with Intrada Amino Acid column (2.0 x 150mm, 3 µm). Mobile phases were 0.3% formic acid in acetonitrile (A) and 0.08M ammonium formate in 20% Acetonitrile (B). The total flow rate was 0.35 mL/min. The optimized LC gradient program for quantitation was started with 20% B, held for 5 min then increased to 100% B at 14 min, hold for 2 min and returned to 20% B at 16.1 min. The LC column was then conditioned for another 3.9 min resulting in a total run time of 20 min. Injection volume is 5 µL, oven temperature is 35 °C. Mass spectral data were acquired in positive electrospray ionization mode, capillary voltage was set at 2.5 kV, source temperature was 100 °C, cone gas and desolvation gas were set at 20 and 400, respectively. LC-MS analysis for intracellular metabolites were performed on a Waters Acquity UPLC & Synapt G2-S system equipped with Waters Acquity UPLC HSS T3 column (2.1 x 150mm, 1.8 µm). Mobile phases were 0.01 M tributylamine/0.02 M acetic acid in water (A) and MeOH (B). The total flow rate was 0.3 mL/min. The optimized LC gradient program for quantitation was started with 2% B, held for 5 min then increased to 80% B at 30 min, hold for 2 min and returned to 2% B at 32.1 min. The LC column was then conditioned for another 3.9 min resulting in a total run time of 36 min. Injection volume is 5 µL, oven temperature is 40 °C. Mass spectral data were acquired in negative electrospray ionization mode, capillary voltage was set at 2.5 kV, source temperature was 120 °C, cone gas and desolvation gas were set at 20 and 400 L/h, respectively. All the acquisition and analysis of data were controlled by Waters MassLynx 4.1 software and raw data were processed using Quanlynx 4.1 software (Waters Corp.).

### Development of BtaSBML2986

The 1.12.0 version of the Human1 model (Robinson et al., 2020) was used as the template model (Fig. 3). The SBML file of the model and the corresponding genes.tsv file were extracted. The genes listed in the genes.tsv file were annotated with UniProt IDs, and protein sequences encoded by the genes were obtained through these IDs and additional processing steps. The ARS-UCD2.0 genome assembly data (https://www.ncbi.nlm.nih.gov/datasets/genome/GCF_002263795.3/) in GenBank format was retrieved from the NCBI database, and the coding sequences were extracted. Bidirectional BLASTP for homology analysis was performed using DIAMOND (Buchfink et al., 2021).

Following the homology analysis, an ortholog GEM was constructed by mapping the genes in Human1 to their corresponding *B. taurus* genes identified through bidirectional BLASTP. Next, *B. taurus*-specific biological data were extracted from KEGG and Rhea (Bansal et al., 2022; Kanehisa & Goto, 2000). In KEGG, the organism code ‘bta’ for *B. taurus* was used to obtain species-specific metabolic data. Additionally, cattle metabolic information was further refined using Pathway Tools (Karp et al., 2021; Kim et al., 2016). After processing, only the metabolic data linked to Rhea were retained to enhance the robustness of the *B. taurus* GEM, as Rhea is a high-quality resource that provides information not fully available in KEGG.

A draft GEM was constructed by integrating the KEGG and Rhea data. GPRuler (Di Filippo et al., 2021) was then used to reconstruct GPR associations, leading to the final reconstruction of the *B. taurus* GEM, BtaSBML2986. In GPRuler, the code for retrieving biological information from KEGG via application programming interface was modified to address limitations in the amount of data processed in a single request. As KEGG could only handle up to 10 genes per request, gene information was processed in batches of 10 genes each.

When adding new reactions to the model, there is a risk of disrupting stoichiometric consistency. To address this issue, reactions were added individually to the draft GEM, ensuring that unconserved metabolites remained at zero. Reactions that could not be directly incorporated underwent a correction process. This process involved restoring stoichiometric balance by adding or removing water and hydrogen, as these adjustments had minimal impact on the overall model structure. By incorporating the corrected reactions, the number of unconserved metabolites was maintained at zero, ensuring a high consistency score when assessed using MEMOTE (Lieven et al., 2020).

### Validation of BtaSBML2986 by using cell growth and media composition data

BtaSBML2986 was transformed into cell-specific genome-scale metabolic models (GEMs) by integrating RNA-seq data from 72 RNA-seq profiles under different conditions (Fig. 5A). The fast task-driven integrative network inference for tissues (ftINIT) method (Gustafsson et al., 2023) was used to generate conditional GEMs. This transformation allowed for more accurate simulations of cell-specific metabolic behaviors and enabled the assessment of growth rates under the given experimental conditions.

Experimental cell growth rates were determined by using linear Relative Growth Rate (RGR), calculated from the cell number at 72 h relative to the initial cell number of 10,000 cells, due to lack of data points (Lamont et al., 2023). The growth rate was calculated as (cell number at 72 h − 10,000) / (10,000 × 72 h). While additional sampling points would enable a more detailed analysis using exponential or logistic growth models, the current dataset, comprising only two sampling points, necessitated the use of this practical approach for estimating growth dynamics.

For GEM-based growth rate estimation, medium composition analysis was conducted to determine the medium conditions for each context-specific GEM. For analysis of medium components, extracellular amino acids were measured using the HITACHI Amino Acid Analyzer L-8900, and extracellular glucose was analyzed using a Waters Acquity Premier HPLC system equipped with a BIO-RAD HPX-87H column. The analysis accounted for metabolites absorbed over the 72 h cultivation period. FBA was then performed using the biomass equation as the objective function to predict cell growth rates.

### Validation of BtaSBML2986 by using metabolome data

The metabolome dataset included measurements of 70 intracellular metabolites under various experimental conditions during the proliferation phase, encompassing three cell sample sets, two sampling points (P1 and P3), and three media conditions (Fig. 5B). Two samples, ‘No_43_P3_965’ and ‘No_66_P3_965’, were excluded due to infeasible solutions during LAD simulation (Lee et al., 2022). The remaining data were analyzed to identify metabolites with significant differences between P1 and P3 sampling points. For validation with GEMs, a process was implemented to compare significantly different metabolites identified from metabolomic data with those derived from flux-sum calculations in the cell-specific GEMs.

Flux-sum values were determined using LAD-based intracellular flux predictions. Flux-sum is defined to be a total sum of all fluxes associated with the consumption or generation of each metabolite (Lee et al., 2024). In this study, LAD minimizes the Manhattan distance of reference flux weights, calculated from RNA-seq, to predict intracellular fluxes (Lee et al., 2022; Kim et al., 2011). As a result, all the reaction fluxes were predicted in accordance with the reference flux weights obtained using RNA-seq data. The biomass flux was constrained to match experimentally determined growth rates to enhance prediction accuracy.

To remove batch effects and ensure consistency across samples, quantile normalization was applied to both metabolome data and flux-sum values. This method ranks features (metabolites or flux values) across all samples, calculates the median for each rank, and replaces the original values with the corresponding median, ensuring consistent distributions while preserving relative rankings. After normalization, significantly different metabolites between P1 and P3 were identified using the SciPy ranksums function. To get significantly different metabolites, the Mann-Whitney U test was performed with BH-corrected P-value < 0.05. The results from the metabolome data and flux-sum analysis were compared to evaluate consistency and cross-validate significantly abundant metabolites. The model evaluation metricswere calculated as follows: accuracy, 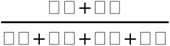; precision, 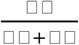, recall, 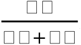, and F1, 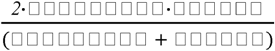.

## Acknowledgments

RNA-seq data and metabolome data for Hanwoo was obtained from the Animal Genomics and Bioinformatics Division at National Institute of Animal Science. This study was supported by the KOBIC Research Support Program. This study was also supported by the ERC Center (RS-2022-NR070840) and Bio & Medical Technology Development Program (RS-2024-00392438) of the National Research Foundation (NRF) funded by Ministry of Science and ICT.

